# Paper based microfluidic aptasensor for food safety

**DOI:** 10.1101/162438

**Authors:** Xuan Weng, Suresh Neethirajan

## Abstract

Food analysis is requiring rapid, accurate, sensitive and cost-effective methods to monitor and guarantee the safety and quality to fulfill the strict food legislation and consumer demands. In our study, a nano-materials enhanced multipurpose paper based microfluidic aptasensor was demonstrated as a sensing tool for accurate detection of food allergens and food toxins. Graphene oxide (GO) and specific aptamer*-*functionalized quantum dots (QDs) were employed as probes, the fluorescence quenching and recovering of the QDs caused by the interaction among GO, aptamer-functionalized QDs and the target protein were investigated to quantitatively analyze the target concentration. The homogenous assay was performed on the paper based microfluidic chip, which significantly decreased the sample and reagent consumptions and reduced the assay time. Egg white lysozyme, ß-conglutin lupine and food toxins, okadaic acid and brevetoxin standard solutions and spiked food samples were successfully assayed by the presented aptasensor. Dual-target assay was completed within 5 min, and superior sensitivities were achieved when testing the samples with commercial ELISA kits side by side.

## 1. Introduction

Food safety has become a critical and global issue for health security and sustainability of both humanity and economy. The growing concern over health risks associated with food allergy, food poisoning and food-borne illness have prompted the evolution of food safety regulation as well as the food analysis methods. Enzyme Linked Immunosorbent Assay (ELISA) (Asensio et al., 2008), PCR (Rodríguez-Lázaro et al., 2013), HPLC (Nollet and Toldrá, 2012), mass spectrometry (Di Stefano et al., 2012), are the most common methods for food analysis to date.

In this study, we aimed at developing a rapid, accurate and cost effective method for food safety monitoring. Two food allergens, egg white lysozyme, ß-conglutin lupine, and two seafood toxins, okadaic acid (OA), brevetoxins, were selected as the testing models. Hen egg is known as one of the most common cause of food allergies both in children and adults. Lysozyme of egg origin is one of the main egg white proteins and being increasingly used in the dairy industry as an antibacterial additive to prevent spoilage of many foodstuffs such as cheese and wine, as well as some medicinal products (Benedé et al., 2014). However, lysozyme is a potential food allergen and accounts for 10-20% of egg allergy which may cause immediate or late adverse reactions such as vomiting, nausea, itching, urticarial and so on (Marseglia et al., 2013). Lupine (Lupin) is a legume belongs to a diverse genus of *Fabaceae* family which is characterized by long flowering spikes (Lupins.org, http://www.lupins.org/lupins/). It has been intensively used in food due to its high value in nutrition and can be found in a wide variety of food products including bread, pasta, sauces, beverages and meat based products such as sausages(ASCIA, http://www.allergy.org.au/health-professionals/papers/lupin-food-allergy).However,lupine allergy is on the rise and hidden lupine allergens in food are a critical problem for lupine sensitive individuals since even very low amounts of lupine may trigger allergic reactions, in severe cases it may lead to life-threatening anaphylaxis (Lupine ELISA Package Insert). Lupine has recently been added to the declaration list of ingredients requiring mandatory indication on the label of foodstuffs within the European Union (Stanojcic-Eminagic, 2010). β-conglutin, one of the two major lupine storage proteins, the other is α-conglutin, accounting for 45% of the total protein content in Lupine is reported to be responsible for lupine allergenicity (Nadal et al., 2012).

Harmful algal bloom (HABs) outbreaks has reportedly intensified throughout the world and pose a grave threat to public health and local economies. HAB toxins through food may cause human diseases by releasing several shellfish toxins, including neurotoxic shellfish poison (NSP), diarrhetic shellfish poison (DSP), paralytic shellfish poison (PSP), ciguatera fish poison (CFP), etc.(Nadal et al., 2014; Christian and Luckas, 2008; Lin et al., 2015) NSP typically affects the gastrointestinal and nervous systems and is caused by consumption of contaminated shellfish with brevetoxins primarily produced by the dinoflagellate (Watkins et al., 2008). Okadaic acid (OA) is a marine toxin, which may cause the diarrheic shellfish poisons (DSP) produced by some unicellular algae from plankton and benthic microalgae (Sassolas et al., 2013a). It is hardly to directly identify OA because OA usually does not affect the smell, appearance and the taste of the seafood (Sassolas et al., 2013b).

At present, there is no treatments available to cure food allergy but only the symptoms after the occurrence of allergic reaction, therefore, food-allergic individuals are typically advised to totally avoid the offending food(s) to protect themselves against a food allergy reaction (Noti et al., 2014), essentially assuming that the threshold dose is zero. In addition, considerable variation of individual threshold dose exist among those with a given type of food allergy (Taylor, et al., 2013). A simple, accurate method for rapid quantitative analysis of allergens or toxins in food is necessary to ensure compliance with food regulations as well as to provide consumer protection.

A multipurpose of PDMS/paper microfluidic aptasensor functionalized by graphene oxide was developed in our study. Aptamers are single-stranded oligonucleotide, or peptide sequences of highly affinity and specificity to various classes of target molecules, which possess many advantages including high sensitivity, specificity and reproducibility as well as the low cost over antibodies thus make themselves be wildly used as the recognition elements in biosensors (Feng et al., 2014). In recent years, graphene oxide (GO) emerges as a good candidate in materials science as well as molecular diagnostic tools due to its unique optical, electronic, thermal, and long-lasting biocompatibility properties (Song et al., 2013). GO has significant and high quenching effect on various fluorophores (Huang and Liu, 2013) via the non-radioactive electronic excitation energy (Swathi and Sebastian, 2009) and its large absorption cross section (Geim and Novoselov, 2007), hence it can be utilized to be quencher in a molecular recognition event.

In this work, we employed GO into the aptasensor and used quantam dots (QDs) as the fluorescence label by considering it has highly chemical stability, efficient and stable fluorescence signals. This was a nano-materials enhanced assay, aptamers specific to the targets, egg white lysozyme, ß-conglutin lupine, okadaic acid and brevetoxins, were firstly bound onto the QDs. After mixing with the GO, the fluorescence was quenched via the process of Förster resonance energy transfer (FRET). In the presence of target proteins in the food sample, the quenched fluorescence would be recovered, the intensity of which was dependent on the concentrations of the target proteins in the food samples. The multipurpose of PDMS/paper microfluidic platform was able to do a dual-target detection with a significantly reduced sample volume with a short time. The use of porous paper as the substrate support for specific aptamer bound QDs-GO probes can effectively simplify the procedures and reduce the cost, because no surface modification is required. Standard solutions were assayed to create the standard curves. Afterwards, a series of food samples were assayed side by side on the presented microfluidic aptasensor and the commercial ELISA kits to evaluate its performance. The results by the aptasensor highly agreed with those from the ELISA kits while significantly reducing the sample and reagent consumptions and having superior sensitivities.

## 2. Experimental

### 2.1. Materials and reagents

The design of the aptamers specific to target analytes, namely egg white lysozyme, ß-conglutin lupine, okadaic acid and brevetoxin, were selected by referring to previous studies (Nadal et al., 2012; Tran et al., 2010; Eissa et al., 2015; Gu et al., 2016) and synthesized by IDT technologies (Coralville, Iowa, USA), the sequences of the selected aptamers are listed in Table 1, all of which were modified with biotin at the 5’end.

**Table 1.**
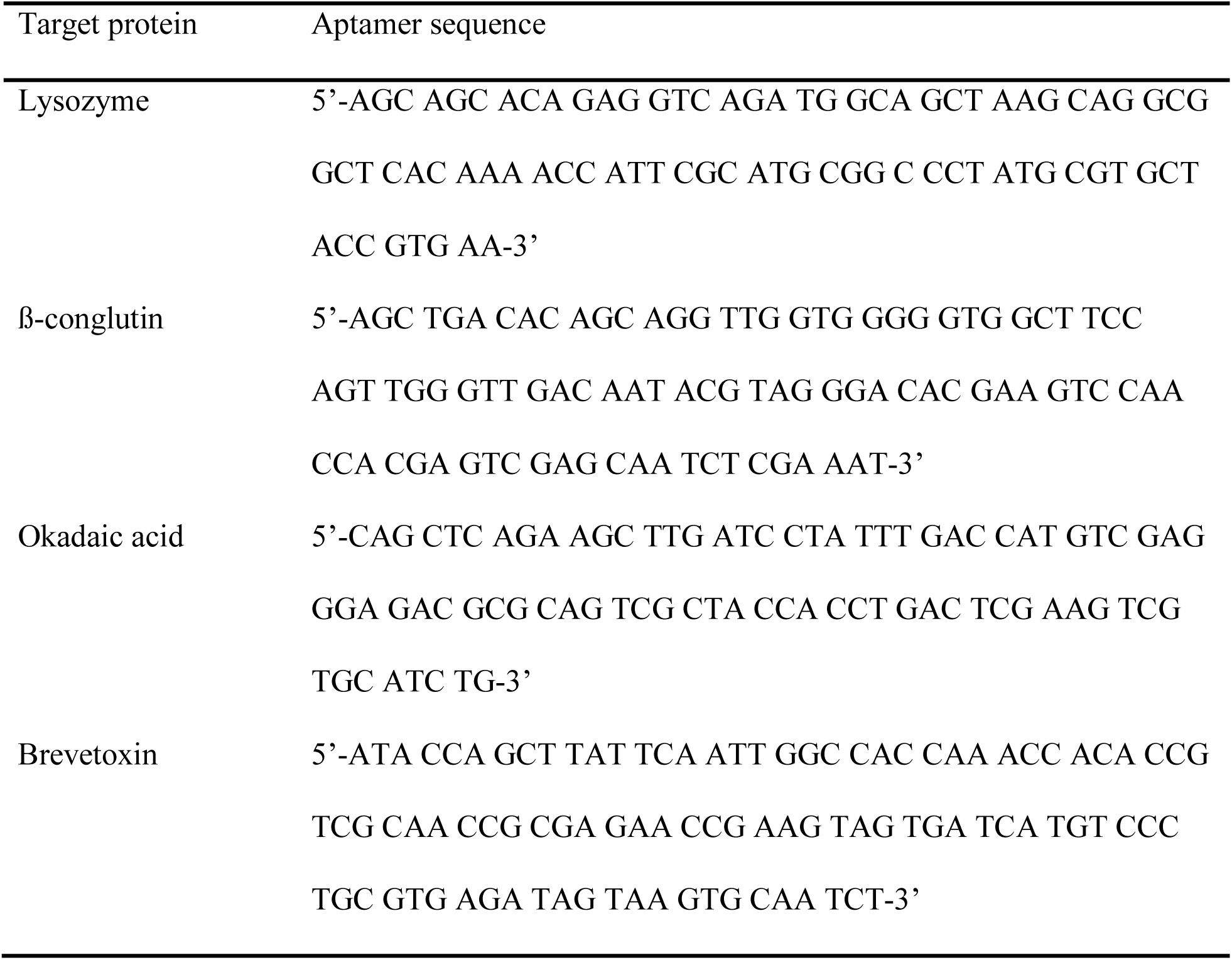
Sequences of selected aptamers

Food Lupine ELISA Test Kit and Brevetoxin (NSP) ELISA Kit were purchased from Creative Diagnostics (Shirley, NY, USA), Lysozyme ELISA Kit and Okadaic Acid (DSP) ELISA Test Kit were obtained from LifeSpan BioSciences, Inc (Seattle WA, USA) and Bioo Scientific Corporation (Austin, Texas, USA), respectively. CdSe Quantum dots modified with covalently attached streptavidin (Qdot^®^ 545 ITK™ Streptavidin Conjugates) were purchased from Invitrogen Life Technologies (Burlington, ON, Canada). Polydimethylsiloxane (PDMS, Sylgard,184) was obtained from Dow Corning (Midland, MI, USA), SU-8 photoresist and developer were obtained from MicroChem Corp. (Westborough, MA, USA). Whatman chromatography paper, graphene oxide, phosphate-buffered saline (PBS), bovine serum albumin (BSA), methanol and all other mentioned chemicals and reagents were purchased from Sigma-Aldrich (Oakville, ON, Canada). Unless otherwise noted, all solutions were prepared with ultrapure Milli-Q water (18.2 MV cm). Eggs, mussels and all other food samples were purchased from grocery stores in Guelph (ON, Canada).

### 2.2. Preparation of aptamer-QDs functionalized GO

The detailed preparation and optimization of aptamer-QDs functionalized GO can be found in our previous study (Weng and Neethirajan, 2016). Briefly, biotinylated aptamer of 10 μM in folding buffer (1 mM MgCl_2_, 1×PBS, pH 7.4) was heated at 85°C for 5 min. The cooling down aptamer was mixed with streptavidin-conjugated quantum dots of 2 μM to proceed with the covalent linking via streptavidin-biotin interaction. The mixture was then brought to 200 μL with PBS and gently shaken for 12 hours under room temperature (RT) condition. The aptamer-QDs conjugates was then obtained by subjecting to ultrafiltration (Amicon Ultra-0.5 mL centrifugal filters, MWCO 50 kDa, EMD Millipore Inc.) with PBS at 6000 rpm for 15 min, repeated three times. The purified aptamer-QDs conjugates re-suspended in 500 μL of PBS and kept at dark for further use.

### 2.3. Microfluidic biochip fabrication and signal capture

The schematic of the paper/PDMS microfluidic chip and the pictures of the real chip are shown in Fig. 1. A high resolution transparency photomask bearing microchannel layout design was firstly drawn by AutoCAD software and printed by Fineline Imaging (Colorado Springs, CO, USA). A master mold was then prepared using 2025 negative photoresist SU-8 by standard photolithography. A thin layer of SU-8 was spin-coated on the surface of the wafer, followed by prebaking at 65°C for3 min and 95°C for 9 min on a hotplate. Afterwards, the photomask was placed onto the coated silicon wafer and exposed to UV using a UV exposure system (UV-KUB, Kloé, France). A mater mold was ready after the post-baking, development and hard-baking. The simple PDMS/paper microfluidic chip consisted of two PDMS layers and a glass slide. The bottom layer of PDMS carried two pairs of wells (□=3mm) for housing well-cut chromatography paper (□=3mm) with QDs-aptamer-GO coating. These dimensions were obtained after calculation by considering many factors, for example, the volume of the sample loading well has to be sufficient to fill out all the four reaction well but not overflow to the waste wells. The two pairs of reaction wells were designed for dual-target detection with duplicate readouts to reduce the testing error. The reaction wells were also the detection wells for fluorescence signal measurement. The top layer of PDMS bearing sample inlet, outlets and associated dispensing channels. The sample dispensing channel is 200 μm in width and 80 μm in depth. The outlets were designed on the top layer of the PDMS so that it may keep the solution stay in the reaction well instead of flowing towards the outlets. Both of the PDMS slabs were created by following the standard soft lithography protocol. Briefly, a mixture of prepolymers of PDMS (10:1 w/w ratio of PDMS and curing agent) was poured onto the master mold at 75°C for 4 h after degassing. The bottom layer of PDMS slab carrying four reaction wells was punched and bond onto the glass. The wells were filled with 0.1% BSA (w/w) for 10 min, washed with 1× PBS and left to dry at RT to reduce the non-specific adsorption of proteins of the PDMS wall (Windvoel et al., 2010). The top layer of PDMS slab was also be punched to form the inlet and outlets. Afterwards, both of these two components were undergone the plasma treatment for bonding and the chromatography paper adsorbing specific aptamer bound QDs-GO probes was placed into the wells before bonding. Then a paper/PDMS microfluidic chip was ready for use.

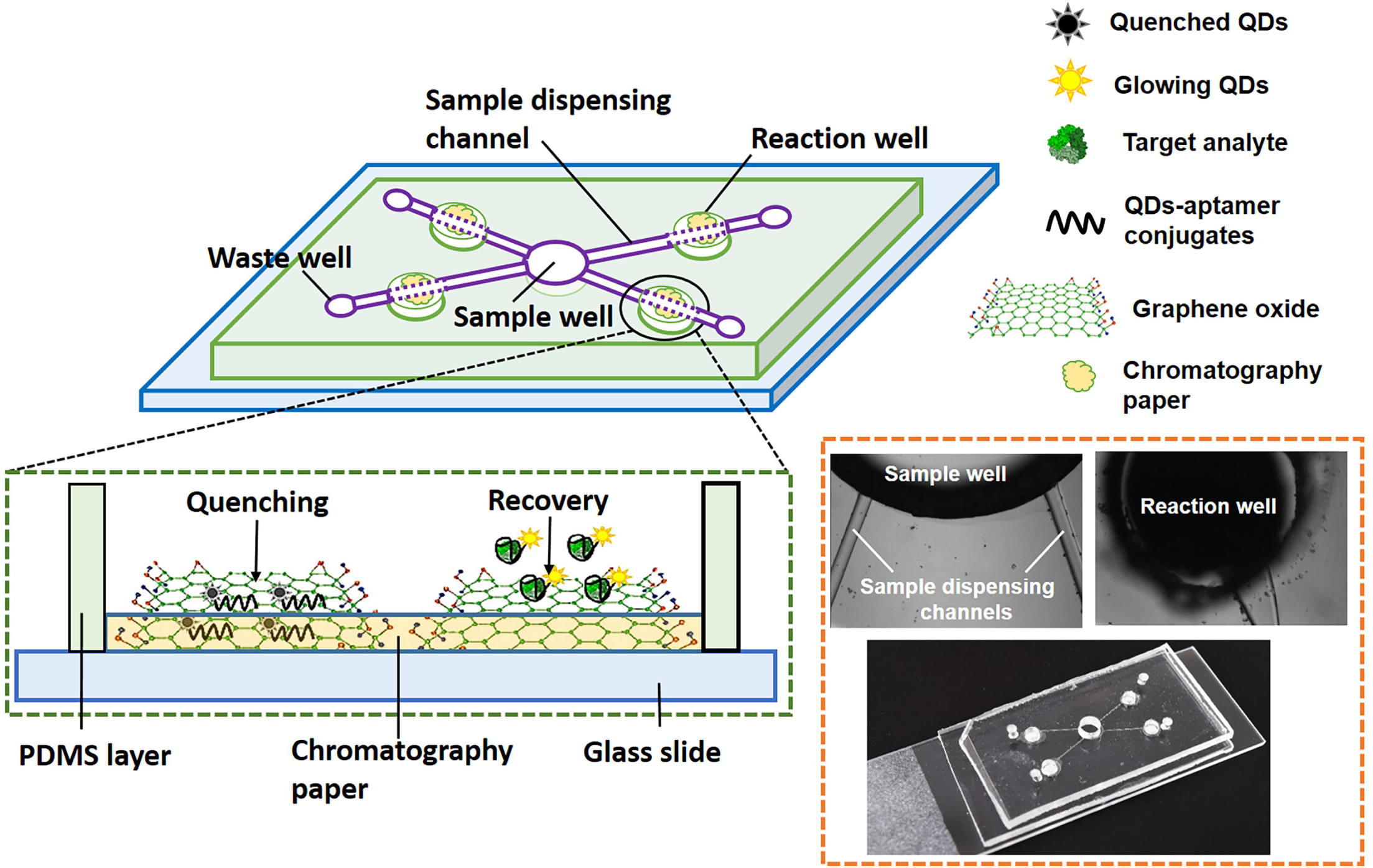

### 2.4. Preparation of food sample

Food sample preparation was conducted by following the procedure indicated in the manual of the commercial kits. Briefly, fresh egg white were firstly diluted with sample diluent to make the dilution series (up to 20000-fold) and followed by centrifuging at 4000×g for 10 minutes at 4°C to remove the particulates. The supernatant was then used in the assay. Mussel tissue was taken off the shells, washed by DI water drained the excess liquid followed by homogenization. 0.5 g of homogenized mussel tissue was carefully weighed and added with 2 mL of 50% methonal followed by vortex for 5 min. The mixture was centrifuged at 4000 rpm for 10 min and 0.5 mL of the supernatant was transferred to a new tube, heated at 75°C for 5 min and followed by centrifugation again for another 10 min at 4000 rpm. Then 50 μL of final supernatant was ready for use after addition with 950 μL of 1× Sample Extraction Buffer. Sausage sample were firstly well grinded and 1 g of the homogenized sausage was suspended in 20 mL of pre-diluted extraction and sample dilution buffer followed by 15 min of incubation in a water bath at 60°C with frequent shaking. Afterwards, the sample was centrifuged at 2000 g for 10 min, the supernatant was ready for assay.

### 2.5. Assay procedure

Fluorescence images were taken by a Nikon DS-QiMc microscope camera mounted on the fluorescent microscopy followed by the fluorescent intensity measurement by the Nikon NIS Elements BR version 4.13 software (Nikon Eclipse Ti, Nikon Canada Inc., Mississauga, ON, Canada). All images were taken under the same settings, namely exposure time, magnification,etc. Food sample detection by commercial kits was conducted by following standard ELISA procedure described in the manual.

## 3. Result and discussion

### 3.1. Characterization and Validation

The characterization of the aptamer-QDs by dynamic light scattering (DLS) analysis and fluorescence spectra measurement were performed and investigated to confirm the conjugations. The detailed procedures can be found in our previous study (Weng and Neethirajan, 2016). The morphology of the graphene oxide and QDs were characterized by TEM imaging as shown in Fig. 2. The hydration diameters of the QDs before and after conjugation (lysozyme aptamer) were measured and compared by dynamic light scattering (DLS) analysis. As shown in Fig. 2(A), the mean hydration diameter of the QDs increased, which verdicted the successful conjugations. Before the on-chip test, the standard solution of these four analytes were measured on the Cytation 5 Multi-mode Reader (BioTek, Winooski, VT, USA) to validate the occurrence of sensing events. The results are shown in the Fig. 3, differentiable fluorescence spectra dependent on the sample concentrations were observed.

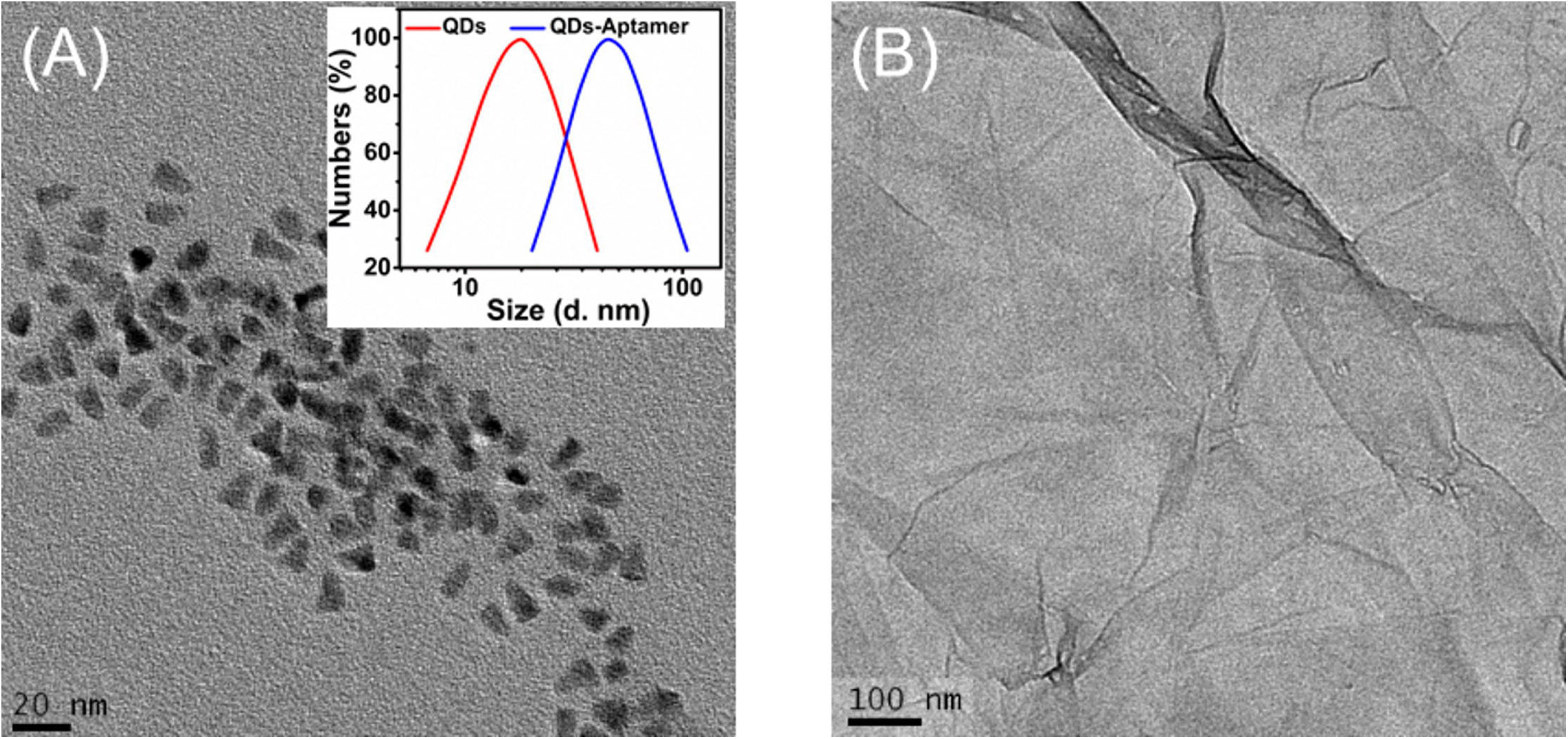

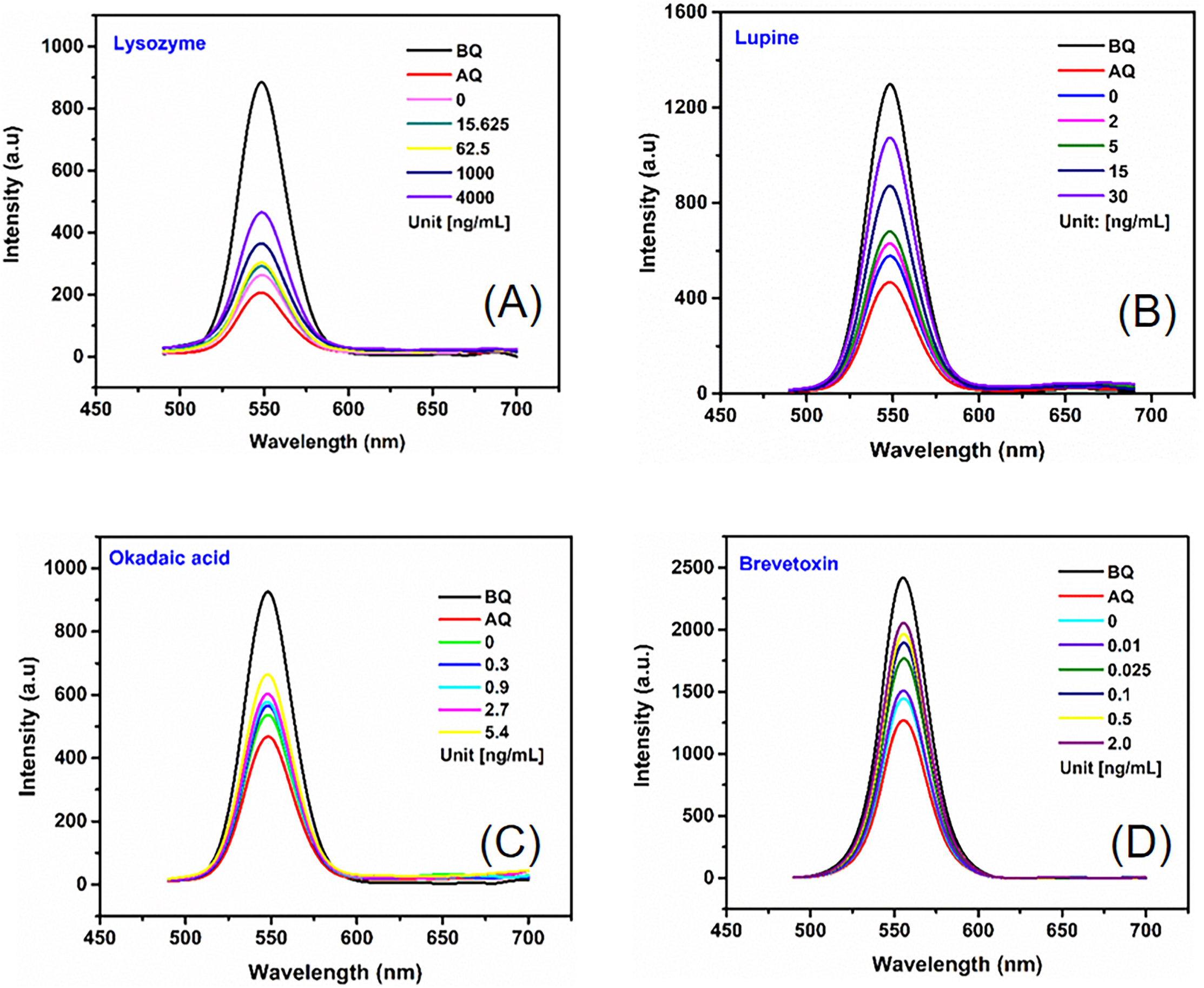

### 3.2. On-chip test

Ten microliters of standard solutions or samples were loading into the central well of the microfluidic chip and dispensed into the four reaction wells by capillary force and wet the chromatography paper with modification of GO-aptamer-QDs. The fluorescence intensities after quenching and recovery were scanned and recorded by the fluorescence microscope, the intensity differences in between were employed to determine the concentrations of the target. Pieces of chromatography paper with the same aptamar-specific GO-QDs were placed in two wells for a duplication. Hence the designed PDMS/paper microfluidic chip was able to achieve the dual-target detection with duplication. Fig. 4 gives an example of the fluorescence images taken before quenching (BQ), after quenching (AQ) and after recovery (RC) by assaying egg white lysozyme of various concentrations. The mean fluorescence intensity of the overall of the reaction well was then analyzed via the Nikon NIS Elements BR software.

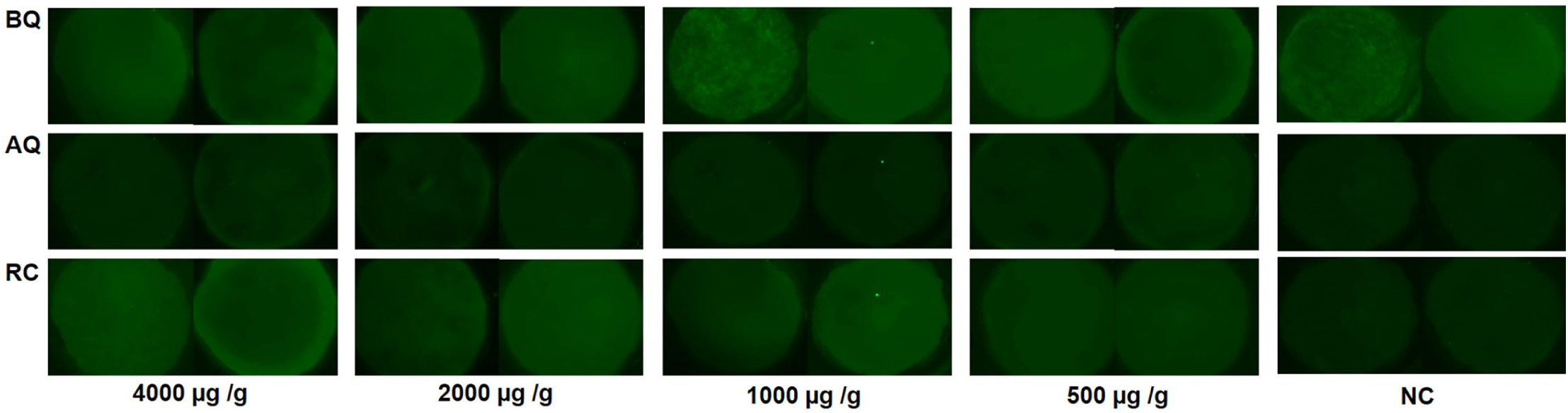

The standard curves were obtained by plotting the mean fluorescence intensity for each stand on the Y-axis against the target concentrations on the X-axis, a linear fit curve were created through the points. The fit curves were presented in the plots shown in Fig. 5, the linear regressions of 0.9469, 0.9839, 0.9838 and 0.975 were calculated and obtained for egg white lysozyme, ß-conglutin lupine and food toxins, okadaic acid and brevetoxin-2 standard solutions, respectively. The calculated limits of detection (Thomsen et al., 2003) based on the standard curves are 343 ng/mL, 2.5 ng/mL, 0.4 ng/mL and 0.56 ng/mL, respectively. These limits of detection by presented aptasensor are superior or comparable to those claimed by the ELISA kits (16 ng/mL, 30 ng/mL, 200 ng/mL and 0.16 ng/mL).

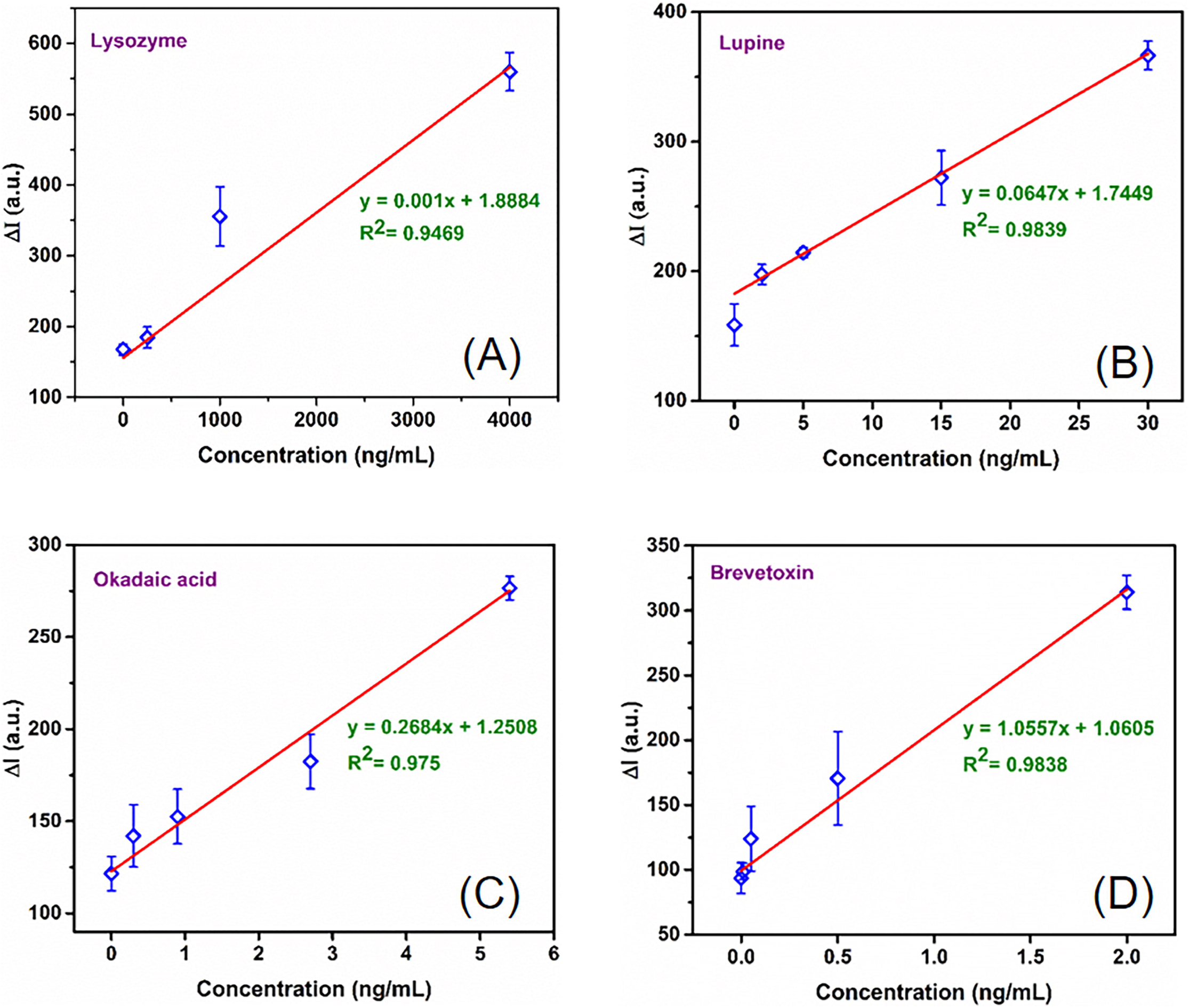

### 3.3. Food samples detection

Spiked food samples were detected by both the on-chip method and the ELISA kits for egg white lysozyme, lupine, okadaic acid and brevetoxin to investigate the accuracy of the on-chip method. Standard solutions were firstly assayed to obtain the standard curves, as shown in Fig. 5. The precision of this method in terms of recovery rate was evaluated by detecting spiked food samples, fresh egg white, mussels, sausages and breads. Each concentration was performed three times to ensure the consistency of the response trend and the recovery rate was calculated as follows:

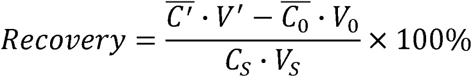

where 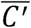 and 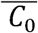 the mean target concentration of the spiked sample and the blank sample, respectively. c_S_ is the concentration of the standard solution spiked into the sample. Vʹ, V_0_, and V_S_ are the volumes of the final spiked sample, blank sample and the standard spiking solution, respectively.

Samples of lysozyme, lupine, okadaic acid and brevetoxin ranging from 0~4000 μg/g, 0~30 μg/g, 0~16.2 μg/g and 0~2 μg/g, respectively, were spiked and tested. The results in Table 2 show the spiked recoveries measured by presented aptasensor were consistent with ELISA kits. As listed in the table, recovery rates of (91.8±2.73)%~(110.18±3.54)%, (89.25±8.30)%~(116.68±10.52)%,(89.63±7.33)%~(105.00±11.46)% and (88.00±9.17)% ~ (112.53±12.22)% were measured in egg white lysozyme, lupine, okadaic acid and brevetoxin for the spiked food samples.

**Table 2.**
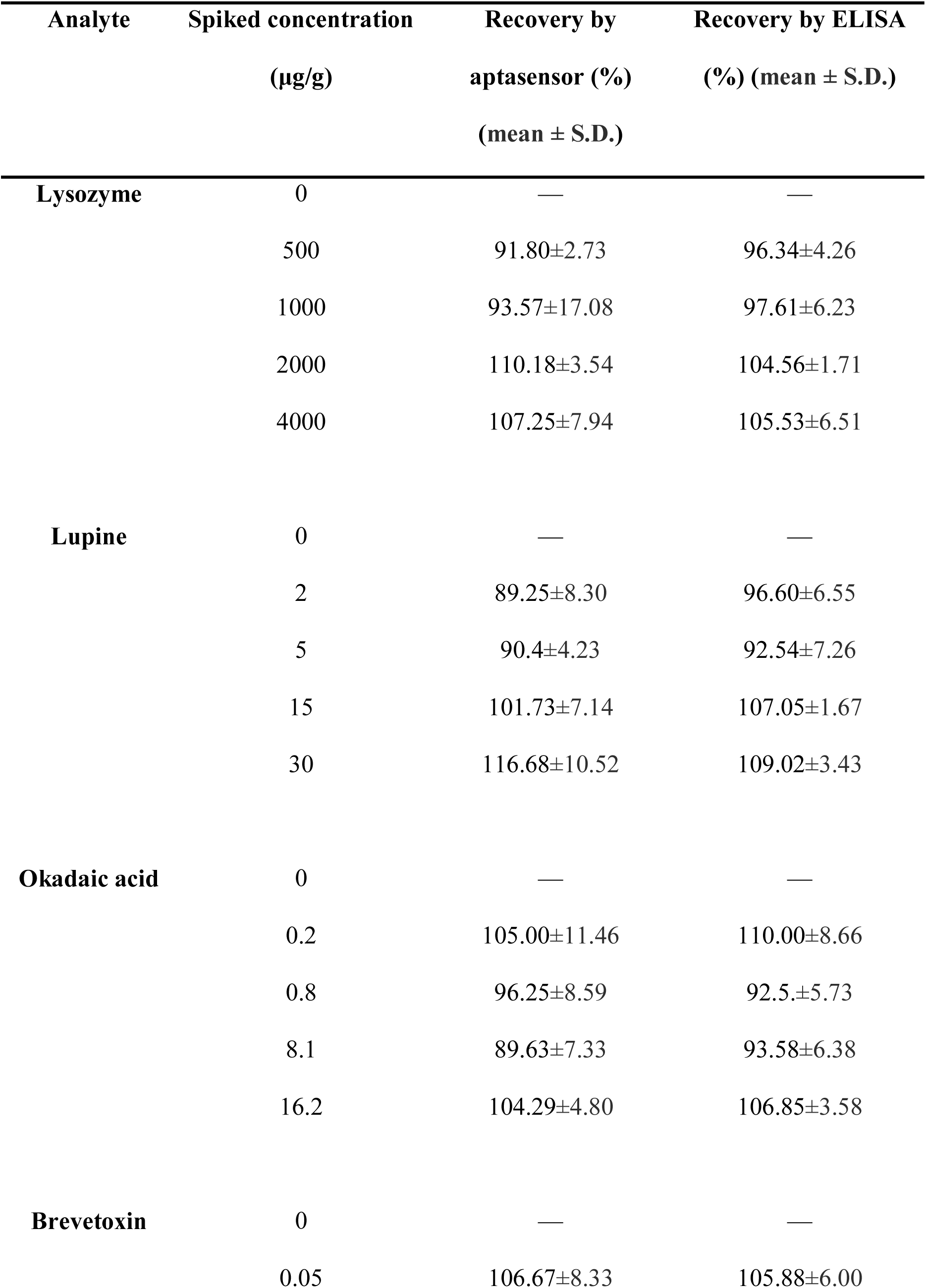
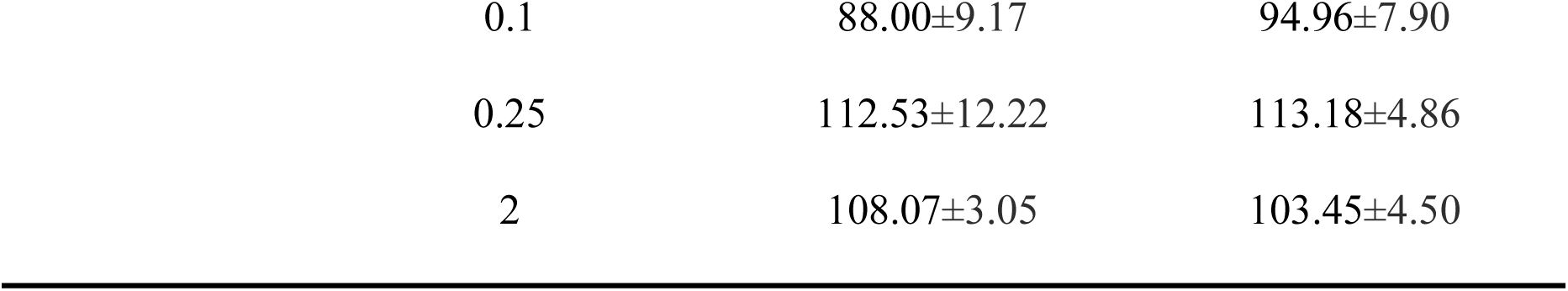
Determination of target analytes concentration in spiked food samples by our aptasensor and ELISA kits

## 4. Conclusions

In this study, a multipurpose PDMS/paper microfluidic aptasensor was used in food analysis, as testing models, food allergens (egg white lysozyme ß-conglutin lupine) and seafood toxins (okadaic acid and brevetoxin-2) were determined in the aptasensor. PDMS is a common used material for fabricating microfluidic device due to its good optical transparency and biocompatibility, non-toxicity and reusability. The utilization of porous paper as the substrate for the specific aptamer bound QDs-GO probes avoids the complicated chemical surface modification thus simplify the whole procedures. The low cost of PDMS and the paper is another advantage of our aptasensor. In addition, the PDMS/paper microfluidic aptasensor utilized grapheme oxide as quencher which can quench the fluorescence of quantum dots conjugated onto the target-specific aptamers. The fluorescence is recovered in the presence of target and its intensity is proportional to the concentration of the target. A significantly decreased sample volume (10 μL) was needed and a dual-target detection with a duplicated results could be achieved in a single test within 5 min to reduce the chance of error. Limit of detection of this sensing platform has been carefully investigated, which are 343 ng/mL, 2.5 ng/mL, 0.4 ng/mL and 0.56 ng/mL with the linear regressions of 0.9469, 0.9839, 0.9838 and 0.975 for egg white lysozyme, ß-conglutin lupine and food toxins, okadaic acid and brevetoxin standard solutions, respectively.

The relative low level of performance is usually provided by the paper-based microfluidic devices. However, the experimental results by this PDMS/paper microfluidic aptasensor demonstrated remarkable sensitivity and selectivity due to the enhancement of nano-materials. Compared to ELISA, which is usually used to detect food allergens and toxins in a centralized lab, our method is rapid, highly sensitive, selective, less expensive, environmentally friendly, and easy to handle. The presented method provides a promising way for the rapid, cost-effective, and accurate determination of food allergens or seafood toxins and also presented its potential of on-site determination capability as well as the flexibility for specifically targeted allergens by selecting corresponding aptamer. With more efforts, an image intensity analyzer may be embedded in this microfluidic aptasensor to build a handheld detection device.

## Acknowledgements

This work was supported by the Natural Sciences and Engineering Research Council of Canada (400705).

